## Supplementary Figures for "Uncovering Transcriptional Dark Matter via Gene Annotation Independent Single-Cell RNA Sequencing Analysis"

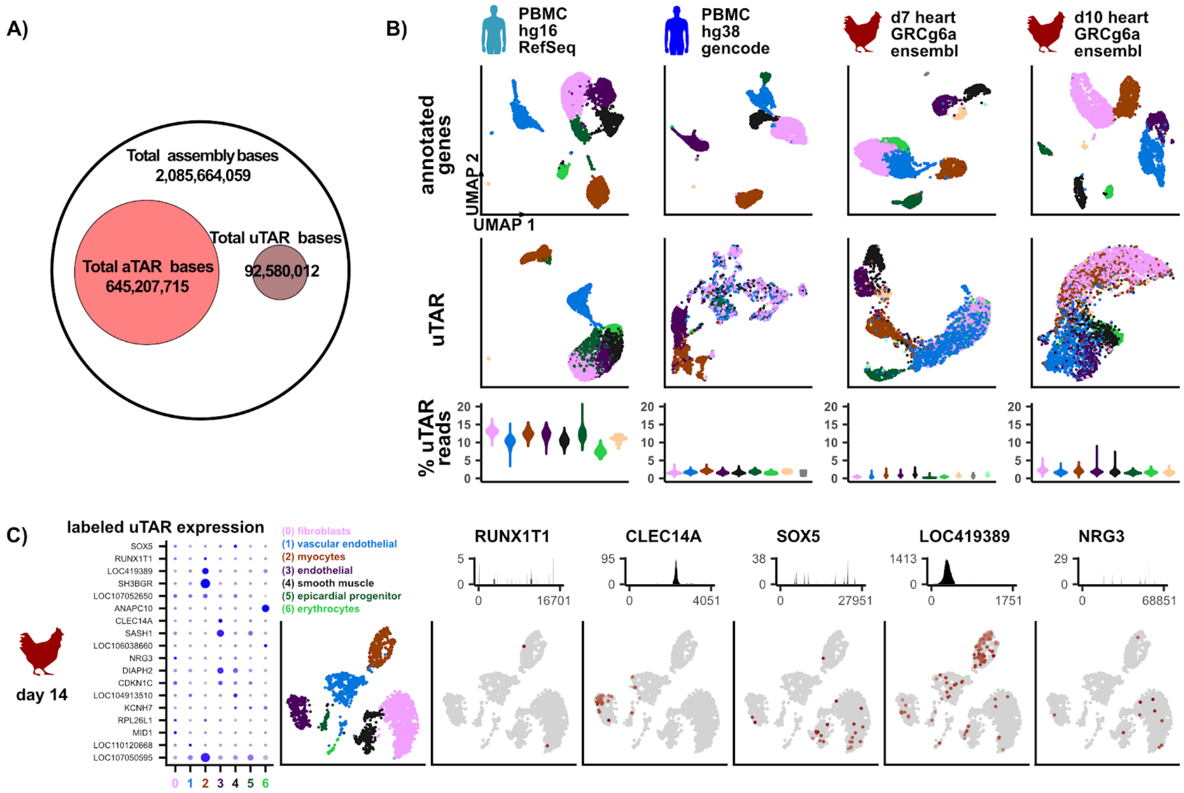


**Extended Data Figure 1 – Additional uTAR analysis and mouse genome (mm10) uTAR statistics. A)** Venn diagram comparing the number of total bases in the mouse (mm10) assembly versus number of bases covered by aTAR and uTAR features. **B)** UMAP dimensional reduction on annotated gene expression features (top row) and uTARs (second row) for human PBMCs relative to hg16 assembly and corresponding RefSeq annotations, human PBMCs relative to hg38 assembly and GENCODE v30 annotations, and in chicken embryonic heart development days 7 and 10. Cells are colored in each column based on gene expression clustering. Relative number of directional uTAR reads for each cell in every cluster also shown as violin plots (third row, colors correspond to UMAPs). 3849 cells in hg16 and hg38, 4132 cells in chicken day 7, and 3313 cells in day 10. **C)** Chicken day 14 heart development dot plot (left) of day 4 differentially expressed uTAR features that are labelled based on sequence homology and cell clusters are numbered along the x-axis. Dot size correspond to number of cells that express the uTAR feature while darker blue color correlate to higher level of log-e normalized expression. UMAP (second left) colored and dimensionally reduced using gene expression features where cell clusters are labeled above the UMAP. Total coverage plot (top) of 5 uTARs along the length of the uTAR feature on the x-axis. The corresponding feature plot on UMAP projection is shown below the coverage plots where darker brown color correlate with higher log-e normalized expression in each cell.

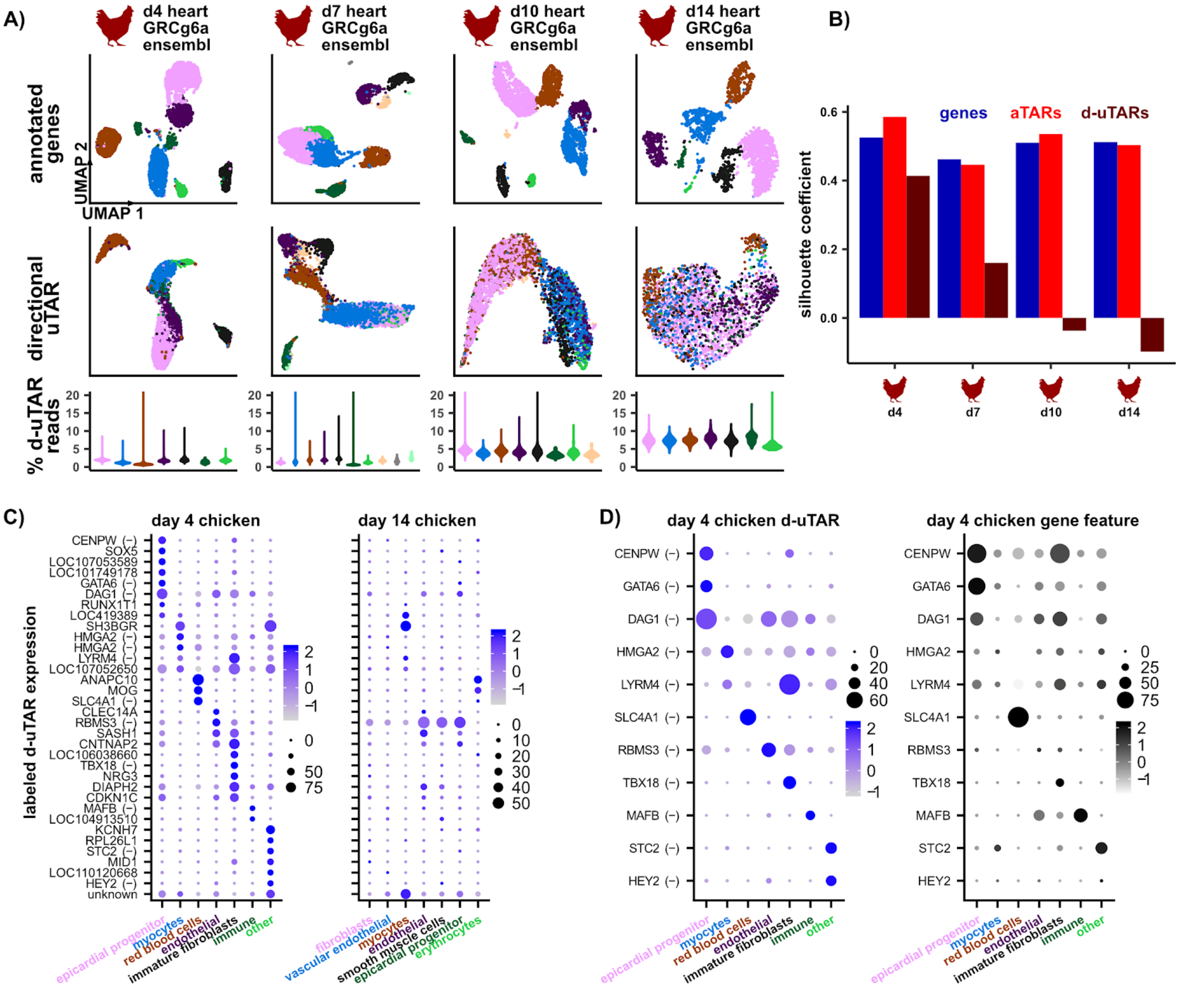


**Extended Data Figure 2 - Accounting for strandedness provides similar results in chicken embryonic heart development. A)** UMAP dimensional reduction on annotated gene expression features (top row) and directional uTARs (second row) for different time points in chicken embryonic heart development. Cells are colored in each column based on gene expression clustering. Relative number of directional uTAR reads for each cell in every cluster also shown as violin plots (third row, colors correspond to UMAPs). 4363 cells in chicken day 4, 4132 cells in day 7, 3313 cells in day 10, and 2198 cells in day 14. **B)** Silhouette coefficient values based on 2D UMAP coordinates of gene expression (blue), aTARs (red), and directional uTARs (maroon) for 4 time points in chicken heart development. Cell labels are defined by gene annotation clustering. **C)** Dot plots of log-e normalized differentially expressed day 4 directional uTAR features in day 4, and 14 labeled based on sequence homology. Features labeled with "(-)" represent directional uTARs found on the antisense strand of the gene. **D)** Dot plot of log-e normalized day 4 directional uTARs on the antisense strand of a gene annotation (left) and the corresponding sense strand gene annotation (right).
